# Unilateral relapse of Behcet’s disease-associated uveitis does not appear to cause asymmetric tear protein profiles

**DOI:** 10.1101/449074

**Authors:** Anyi Liang, Weiwei Qin, Chan Zhao, Youhe Gao

## Abstract

Purpose: To explore whether unilateral relapse of Bechet’s disease uveitis (BDU) causes differences in the tear proteome between the diseased and the contralateral quiescent eye.

Experimental design: To minimize interindividual variations, bilateral tear samples were collected from the same patient (n=15) with unilateral relapse of BDU. A data-independent acquisition (DIA) strategy was used to identify proteins that differed between active and quiescent eyes.

Results: A total of 1,797 confident proteins were identified in the tear samples, of which 371 are also highly expressed in various tissues and organs. Sixty-two (3.5%) proteins differed in terms of expression between tears in active and quiescent eyes, similar to the number of differentially expressed proteins (74, 4.1%) identified in a randomized grouping strategy. Furthermore, the intrapair trend of the differentially expressed proteins was not consistent and none of the proteins showed the same trend in more than 9 pairs of eyes.

Conclusions and clinical relevance: Unilateral relapse of BDU does not appear to cause asymmetric changes in the tear proteome between active and contralateral quiescent eyes. Tear fluid is a valuable source for biomarker studies of systemic diseases.

**Statement of clinical relevance:** Tears are an easily, noninvasively accessible body fluid that is a valuable source of biomarkers for various diseases. Behcet’s disease uveitis (BDU) has high potential to cause blindness and represents the leading cause of morbidity in BD patients, especially in frequently relapsing cases. Here, we adopted a method combining a “dry” method for tear preservation and nano-LC-DIA-MS/MS system to explore whether unilateral relapse of BDU causes differences in the tear proteome between the diseased and the contralateral quiescent eye, with the aim of evaluating tear fluid as a source for biomarker studies of uveitis relapse.

## Introduction

Tear fluid is a complex mixture of water, electrolytes, metabolites, lipids and proteins (mucins, enzymes, glycoproteins, immunoglobulins and others) secreted mainly by the main and accessory lacrimal glands ^[1, 2]^. Tears can be easily and noninvasively accessed ^[3]^ and are considered a useful source for biomarker research of ocular and systemic diseases. According to a recent review, hundreds of potential specific molecular biomarkers in tears were found to be associated with ocular diseases, including dry eye disease, keratoconus and thyroid associated orbitopathy. Other reports showed that tears can reflect the state of breast cancer, prostate cancer and multiple sclerosis ^[2, 4, 5]^. Moreover, tears may reflect central metabolism in some neurological disorders ^[6-9]^.

Behcet’s disease (BD) is a chronic, relapsing systemic inflammatory disease characterized by recurrent oral aphthae and a variety of other clinical features, including uveitis (also known as intraocular inflammation), skin lesions, gastrointestinal involvement, neurologic disease, vascular disease and arthritis ^[10]^.

Half of BD patients may associate with uveitis, which is a common cause of vision loss and blindness worldwide, especially in young men ^[11]^. Interestingly, despite being a symmetrical organ, the eyes of patients with Bechet’s disease uveitis (BDU) with unilateral uveitis relapse show dramatically distinct clinical manifestations. While the quiescent eye appears completely normal, the other active eye might have severe visual acuity impairment possibly caused by anterior uveitis, vitritis, retinal vasculitis, retinitis, cystoid macular edema or papillitis ^[12]^. It is possible that, considering the anatomical proximity, intraocular inflammation may cause substantial changes in the tear proteome through local mechanisms, making tears a potential source for biomarkers of uveitis involving the ipsilateral eye.

To address this hypothesis and to minimize interindividual variations, tear samples were simultaneously collected from both eyes of BDU patients with unilateral uveitis relapse. Samples were then preserved by our previously developed “dry” method ^[13]^ and were further analyzed by a data-independent acquisition (DIA) strategy.

## Materials & Methods

### 1. Patients

BDU patients presenting to our center with acute unilateral relapse of panuveitis between January 2018 and June 2018 were included. The consent procedure and the study protocol were approved by the Institutional Review Board of the Institute of Basic Medical Sciences, Chinese Academy of Medical Sciences. (Project No. 007–2014). Written informed consent was obtained from each subject prior to the study.

Patients were diagnosed with BD according to the criteria of the International Study Group (ISG) or International Criteria for Behcet’s Disease (ICBD) ^[14]^. Acute relapse of panuveitis was defined as a decrease in visual acuity and the presence of a combination of anterior uveitis (more than 0.5 cells in the anterior chamber), vitritis (more than 1 cell), and inflammation of the posterior segment with the presence of at least one of the following: retinal vasculitis, retinitis, cystoid macular edema and papillitis ^[12]^. The contralateral eye was reported to be quiescent for at least two months. Patients with any of the following conditions were excluded: 1) other ocular or orbital diseases, including allergic conjunctivitis, infectious conjunctivitis, keratitis of infectious or noninfectious cause, thyroid associated orbitopathy, primary open angle glaucoma and primary angle closure glaucoma; 2) previous ocular trauma or surgeries; 3) other forms of uveitis; 4) other systemic diseases, such as diabetes, hypertension, cardiovascular diseases, neurological disorders and irrelevant rheumatic diseases; and 5) a Schirmer test of less than 10 mm in either eye.

### 2. Tear sample collection and preservation

Tear samples were collected and preserved using a novel Schirmer strip-based dry method ^[13]^. In detail, Schirmer strips (Tianjin Jingming New Technological Devepoment Co.,Ltd) were placed at the lateral 1/3 of the lower conjunctival sacs of both eyes for 5 min, and strips with tears exceeding 10 mm were collected. The Schirmer strips were dried immediately with a hair dryer (70°C for 1 min) and were stored in properly labeled vacuum-sealed bags at room temperature and transferred to −80°C freezer within 2 weeks.

### 3. Sample preparation for mass spectrometry

Tear protein extraction. The strip was cut into small pieces and transferred to a 0.6 mL tube. Next, 200 mL elution buffer (100 mM NH_3_HCO_3_, 50 mM NaCl) was added and gently shaken for 2 h at room temperature. The tube was punctured at the bottom with a cannula, placed in a larger tube (1.5 mL) and centrifuged at 12,000 g for 5 min ^[15]^.

The filtrate in the outer tube was collected and quantified by the Bradford method. Tryptic digestion. The proteins were digested with trypsin (Promega, USA) using filter-aided sample preparation methods ^[16]^. Briefly, 200 µg of the protein sample was loaded on the 10-kD filter unit (Pall, USA). For digestion, the protein solution was reduced with 4.5 mM DTT for 1 h at 37°C and then alkylated with 10 mM of indoleacetic acid for 30 min at room temperature in the dark. Finally, the proteins were digested with 3 µg of trypsin for 14 h at 37°C. The peptides were digested using Oasis HLB cartridges (Waters, USA). The resulting peptides were dried and desalted in a SpeedVac (Thermo Fisher Scientific, Waltham, MA). The dried peptide samples were resuspended in 0.1% formic acid and were quantified using a PierceTM BCA protein assay kit. Two micrograms of each sample were loaded for LC-MS/MS analysis using a data-independent acquisition (DIA) method.

High-pH fractionation: Equal volumes of tryptic digested peptides of each sample were pooled to generate the pooled sample (100 µg of peptides) for the development of a spectral library and for DIA analysis optimization and quality control. Sixty micrograms of pooled peptides was then fractionated using a high-pH spin column (Thermo Pierce, USA). After equilibration of the column, the dried sample was resuspended in 100% Buffer A (acetonitrile:H_2_O, 90:10 with 0.1% formic acid and 10 mM of ammonium formate) and loaded onto the column, then eluted with Buffer B at 5, 7.5, 10, 12.5, 15, 17.5, 20, 30, 50 and 70% of Buffer B (H_2_O with 0.1% formic acid). Fractionated samples were dried completely and resuspended in 20 μl of 0.1% formic acid. Six microliters of each of the fractions was loaded for LC-MS/MS analysis using a data-dependent acquisition (DDA) method.

### 4. LC-MS/MS setup for data-dependent and data-independent acquisition

LC-MS/MS data acquisition was performed on a Fusion Lumos mass spectrometer (Thermo Fisher Scientific) interfaced with an EASY-nLC 1000 HPLC system (Thermo Fisher Scientific). For both DDA and DIA analyses, the same LC settings were used for retention time stability. For facilitating retention time alignments among samples, a retention time kit (iRT kit from Biognosys, Switzerland) was spiked at a concentration of 1:20 v/v in all samples ^[17]^. The digested peptides were dissolved in 0.1% formic acid and loaded on a trap column (75 µm × 2 cm, 3 µm, C18, 100 A°). The eluent was transferred to a reversed-phase analytical column (50 µm × 500 mm, 2 µm, C18, 100 A°). The eluted gradient was 5–30% buffer B (0.1% formic acid in 99.9% acetonitrile; flow rate of 0.6 μl/min) for 60 min.

To acquire a spectral library for use in DIA data extraction, 6 μl of each of the fractions was analyzed by data-dependent acquisition. The full scan was done with a 60K resolution at 200 m/z from 350 to 1 500 m/z with an AGC target of 1e6 and a max injection time of 50 ms. Monoisotopic masses were then selected for further fragmentation for ions with a 2 to 6 positive charge within a dynamic exclusion range of 30 seconds and a minimum intensity threshold of 1e4 ions. Precursor ions were isolated using the quadrupole with an isolation window of 1.6 m/z. The most intense ions per survey scan (top speed mode) were selected for collision-induced dissociation fragmentation, and the resulting fragments were analyzed in the Orbitrap with the resolution set to 60 000. The normalized collision energy for HCD-MS2 experiments was set to 32%, the AGC target was set at 5e4 and the maximum injection time was set to 30 ms. The DDA cycle was limited to 3 seconds.

For data-independent acquisition, two μg of each sample was analyzed. Survey MS scans were acquired in the Orbitrap using a range of 350–1550 m/z with a resolution of 120 000 at 200 m/z. The AGC target was set at 1e6 with a 50 ms max injection time. Twenty-four optimal acquisition windows covered a mass range from 350 to 1 500 m/z (Supplemental Table S1). The normalized collision energy for HCD-MS2 experiments was set to 32%, the AGC target was set at 2e5 and the maximum injection time was set to 54 ms. A quality control DIA analysis of the pooled sample was inserted after every ten tear samples were tested.

The raw mass spectrometric files were stored at the public repository iProX (URL: http://www.iprox.org/page/PSV023.html;?url=1538466057619tm3u, passwords: IwHL).

### 5. Data analysis

The raw MS data files acquired by the DDA mode for library construction were processed using Proteome Discoverer (version 2.1; Thermo Fisher Scientific, USA) with SEQUEST HT against the SwissProt human database (released in July 2016, containing 20 228 sequences) and the Biognosys iRT peptides sequences. SEQUEST HT Search parameters consisted of the parent ion mass tolerance, 10 ppm; fragment ion mass tolerance, 0.02 Da; fixed modifications, carbamidomethylated cysteine (+58.00 Da); and variable modifications, oxidized methionine (+15.995 Da). Other settings included the default parameters. All identified proteins had an FDR of ≤1%, calculated at the peptide level. To generate the spectral libraries, DDA spectra were analyzed as described above, and a spectral library was generated using the spectral library generation functionality of Spectronaut Pulsar (Biognosys, Switzerland). The library was devised by importing the search results and the raw files using Spectronaut with the default parameters.

The raw DIA files were imported to Spectronaut Pulsar, and the default settings were used for targeted analysis. In brief, a dynamic window for the XIC extraction window and a non-linear iRT calibration strategy were used. Mass calibration was set to local mass calibration. Interference correction on the MS1 and MS2 levels was enabled, removing fragments/isotopes from quantification based on the presence of interfering signals but keeping at least three for quantification. The false discovery rate (FDR) was set to 1% at the peptide precursor level and at 1% at the protein level. The significance criteria for a T-test was a p value <0.01. A minimum of two peptides matched to a protein and a fold change >1.5 were used as the criteria for the identification of differentially expressed proteins.

## Results & Discussion

### 1. Clinical characteristics of BDU patients

A summary of the overall experimental approach is presented in Figure 1. To minimize interindividual variations, bilateral tear samples were collected from the same BDU patient (n=15) with unilateral uveitis relapse. The average age of the patients was 28.6 years old, and the female-to-male ratio was 1:4. Seven and eight patients had active inflammation in the left or right eye, respectively. The volume of each tear sample collected by the Schirmer strips was between 10 and 30 mm (Table 1).

**Figure 1.**
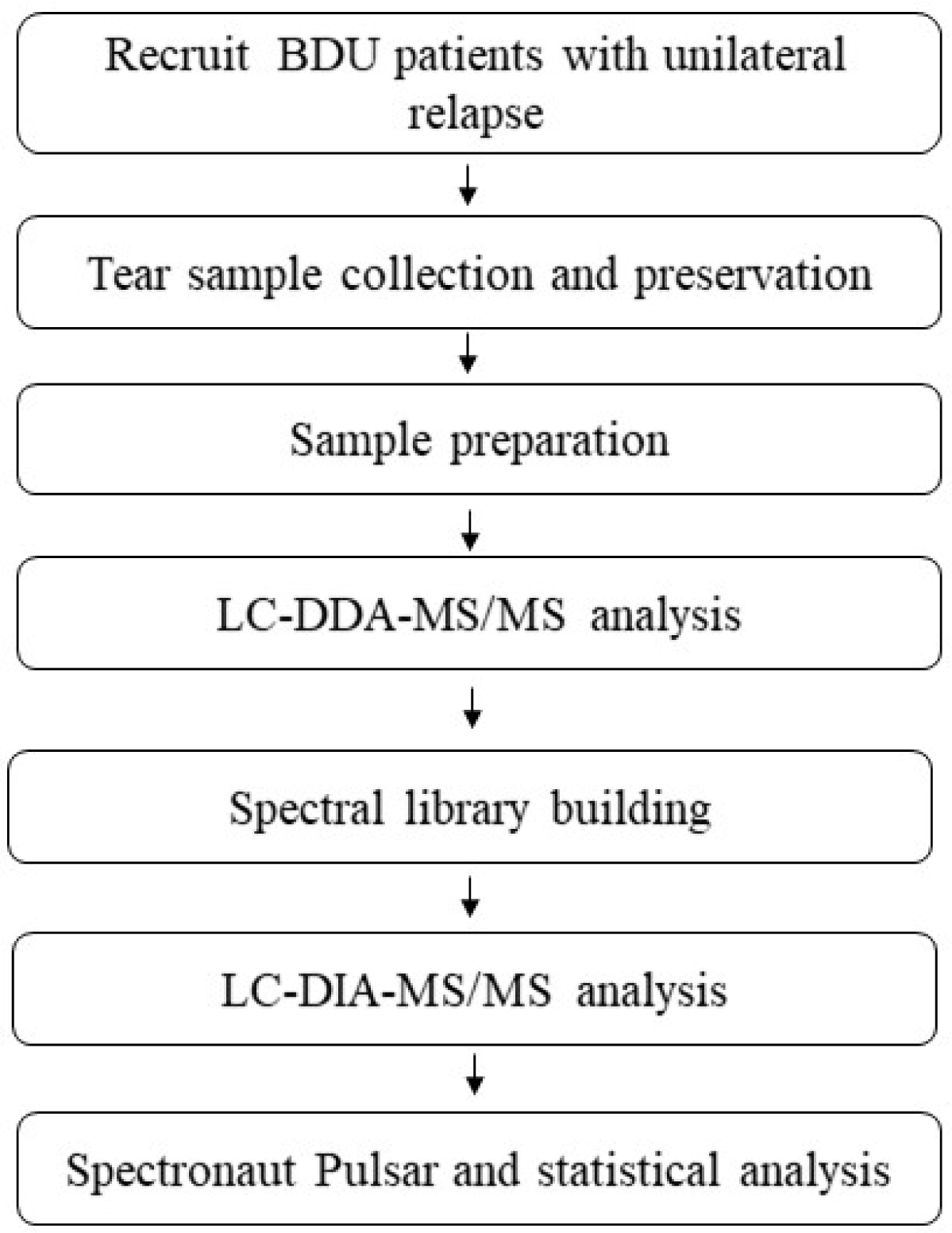
Workflow schematic of this study.

**Table 1.** Demographic characteristics, inflammatory status and volume of tears collected from each eye of the enrolled BDU patients

| Number | Gender | Age | Right eye |  | Left eye |  |
| --- | --- | --- | --- | --- | --- | --- |
|  |  |  | Status | Volume (mm) | Status | Volume (mm) |
| **1** | M | 23 | A | 20 | Q | 30 |
| **2** | M | 30 | Q | 22.5 | A | 30 |
| **3** | F | 26 | Q | 15 | A | 30 |
| **4** | M | 28 | Q | 25 | A | 25 |
| **5** | M | 26 | Q | 10 | Q | 12.5 |
| **6** | F | 32 | A | 12 | Q | 14 |
| **7** | M | 18 | A | 25 | Q | 25 |
| **8** | M | 28 | A | 15 | Q | 20 |
| **9** | F | 29 | A | 30 | A | 25 |
| **10** | M | 28 | A | 10 | Q | 15 |
| **11** | M | 22 | Q | 12 | A | 13 |
| **12** | M | 34 | Q | 23 | A | 20 |
| **13** | M | 25 | A | 17 | Q | 23 |
| **14** | M | 33 | A | 27 | Q | 20 |
| **15** | M | 48 | Q | 18 | A | 22 |

A: active (acute relapse of panuveitis); Q: quiescent (healthy for at least two months).

### 2. Spectral library establishment and DIA method optimization

The spectral library that was generated, as described in the methods, contained 16 135 peptides corresponding to 2 779 protein groups (Supplemental Table S2). Prior to individual sample analysis, the pooled peptide sample was subjected to DIA experiments in order to refine the acquisition windows list and the cycle time (data points per peak). Ultimately, 24 variable windows were used (Supplemental Table S1), resulting in 7–8 data points per peak (cycle time of 1.8 s). The DIA method was benchmarked based on the number of peptide identifications (at 1% peptide and protein FDR) with a CV of MS2 quantification below 20% over triplicate acquisition of the pooled sample.

In total, 1 797 confident proteins were identified in the 30 tear samples from both eyes of the 15 patients. We compared each protein with the tissue-enriched proteome ^[18]^. In all, 371 proteins that were highly expressed in various tissues and organs were also identified in the tear samples ^[13, 18]^. This number is comparable to values from our previous study that identified 365 out of 514 proteins. These results suggest that the tear proteome may reflect changes in a variety of tissues/organs and that tears may be a window for disease biomarker discovery.

### 3. Tear proteome changes

All tear samples were assigned to either an active (A) or a quiescent (Q) group according to the inflammatory status of the eye (Table 1). Differential proteins were screened with the following criteria: fold change ≥ 1.5 between the two groups and a P-value of paired t-test < 0.05. A total of 62 different proteins were identified (details are described in Supplemental Table S3), accounting for approximately 3.5% of the total identified proteins. Considering interindividual variations in tear protein profiles, abundances of the 62 different proteins were compared independently between the eyes of each patient. The trends in abundances of the differentially expressed proteins between the active and the contralateral quiescent eyes were not consistent, and no protein showed the same trend in more than 9 pairs of eyes (Table 2).

**Table 2.** Intrapair trends in the abundance of the identified differential proteins

| Number of sample pairs | Number of proteins* |  |
| --- | --- | --- |
|  | Increased | Decreased |
| **≥10** | 0 | 0 |
| **9** | 3 | 0 |
| **8** | 8 | 2 |
| **7** | 24 | 22 |
| **6** | 29 | 9 |
| **5** | 20 | 28 |
| **4** | 6 | 32 |
| **3** | 2 | 15 |
| **2** | 0 | 4 |
| **1** | 0 | 0 |

\* Number of proteins that had the same intrapair trend (active versus quiescent).

For further verification, an unsupervised cluster analysis was performed. As indicated in Figure 2, the active group was not distinguishable from the quiescent group. When the tear samples were randomized into two groups, irrespective of inflammation status, a similar level of different proteins (74, 4.1%) was detected (Supplemental Table S4). These results suggested that an acute attack of unilateral BDU may not cause significant differences in the tear proteome between active and quiescent eyes. It is possible that unilateral activation of BDU may cause symmetric changes of the tear proteome in both eyes by altering the serum levels of a variety of proteins. As severe as BDU may be, intraocular inflammation may not significantly influence the behavior of the lacrimal glands and other tear-protein secretory cells.

**Figure 2.**
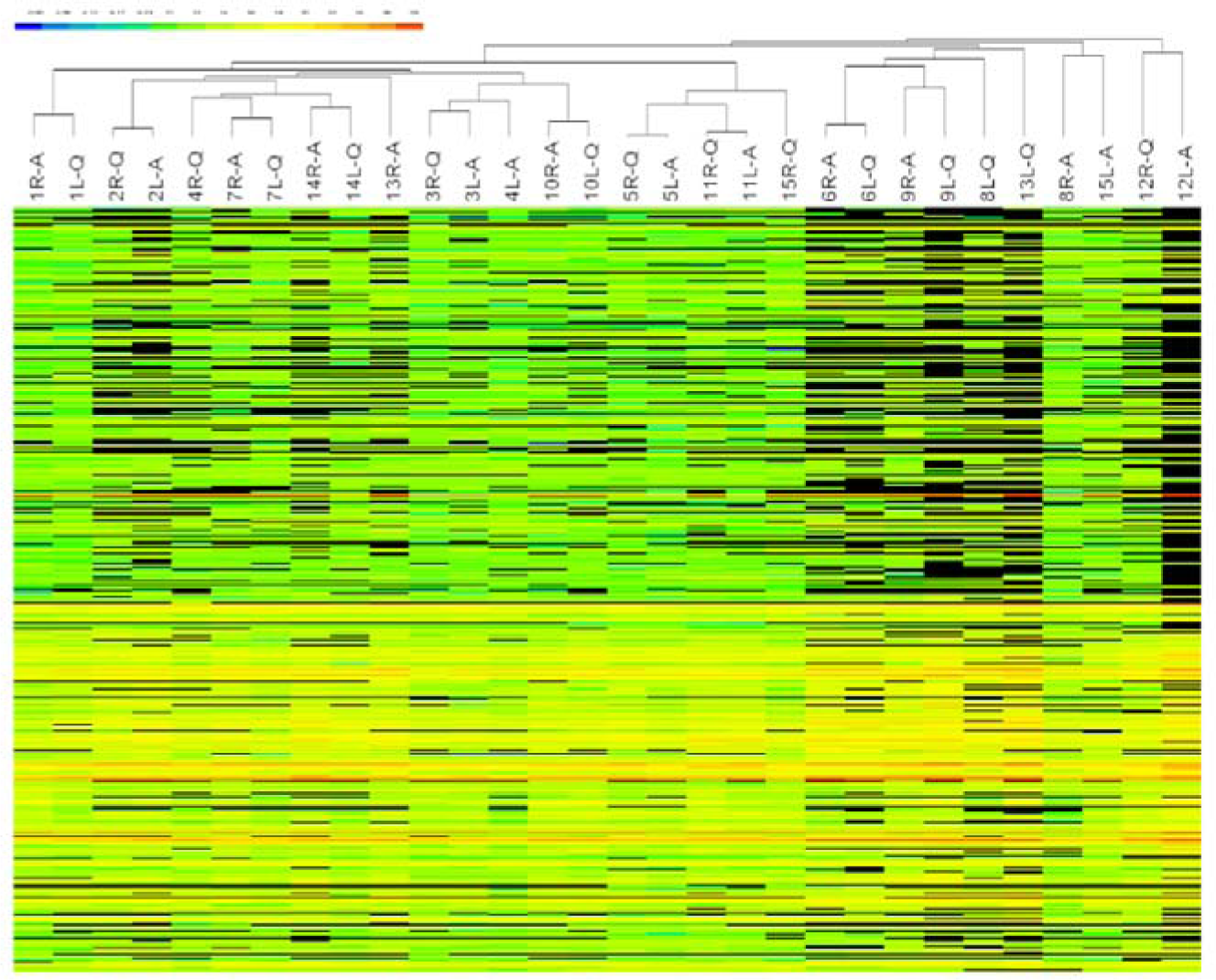
Heatmap of Hierarchical clustering. R: right eye; L: left eye; A: active (acute relapse of panuveitis); Q: quiescent (healthy for at least two months).

Our study was limited by a lack of tear samples from healthy controls or BDU patients in an inactive disease state. In this scenario, comparing tear proteomes between BDU patients and healthy controls or between active and quiescent phases of BDU might lead to discovery of BDU-associated tear biomarkers. While this study was limited by its narrow scope, it provides insight into future tear proteomic studies of intraocular inflammation.

## Conclusion

In BDU patients, unilateral intraocular inflammation did not appear to cause asymmetric tear proteome changes between the active eye and the contralateral quiescent eye. Tear fluid maybe a valuable source for biomarker studies of systemic diseases.

## Acknowledgements

The National Key Research and Development Program of China (SQ2018YFC090062, 2016YFC1306300); Beijing Natural Science Foundation (7172076); the Fundamental Research Funds for the Central Universities (2015KJJCB21); Beijing cooperative construction project (110651103); Beijing Normal University (11100704); and the 2016 PUMCH Science Fund for Junior Faculty (pumch-2016-2.27).

